# Dissociating cholinergic influence on alertness and temporal attention in primates in a simple reaction time paradigm

**DOI:** 10.1101/765883

**Authors:** Vilmos Oláh, Balázs Knakker, Attila Trunk, Balázs Lendvai, István Hernádi

**Affiliations:** Grastyán Translational Research Center, University of Pécs & Gedeon Richter Plc., 6 Ifjúság út, H-7624, Pécs, Hungary; Department of Experimental Zoology and Neurobiology, Faculty of Sciences, University of Pécs, 6 Ifjúság út, H-7624, Pécs, Hungary; Szentágothai Research Center, Center for Neuroscience, University of Pécs, 20 Ifjúság út, H-7624, Pécs, Hungary; Department of Pharmacology and Drug Safety Research, Gedeon Richter Plc., Gyömrői út 19-21, H-1103, Budapest, Hungary; Institute of Physiology, Medical School, University of Pécs, 12 Szigeti út, H-7624, Pécs, Hungary

**Keywords:** Foreperiod, temporal expectation, scopolamine, donepezil, rhesus monkey

## Abstract

The ability to promptly respond to behaviourally relevant events depends on both general alertness and phasic changes in attentional state driven by temporal expectations. Using a variable foreperiod simple reaction time (RT) task in four adult male rhesus macaques, we investigated the role of the cholinergic system in alertness and temporal expectation. Foreperiod-effects on RT reflect temporal expectation, while alertness is quantified as overall response speed. We measured these RT parameters under vehicle treatment and systemic administration of the muscarinic receptor antagonist scopolamine. We also investigated whether and to what extent the effects of scopolamine were reversed by donepezil, a cholinesterase inhibitor widely used for the treatment of dementia. In the control condition, RT showed a continuous decrease as the foreperiod duration increased, which clearly indicated the effect of temporal expectation on RT. This foreperiod effect was mainly detectable on the faster tail of the RT distribution and was eliminated by scopolamine. Furthermore, scopolamine treatment slowed down the average RT. Donepezil treatment was efficient on the slower tail of the RT distribution and improved scopolamine-induced impairments only on the average RT reflecting a general beneficial effect on alertness without any improvement in temporal expectation. The present results highlight the role of the cholinergic system in temporal expectation and alertness in primates and help delineate the efficacy and scope of donepezil and other cholinomimetic agents as cognitive enhancers in present and future clinical practice.

## 1. Introduction

In a simple reaction time (RT) task, subjects have to detect and give speeded response to target stimuli that are either preceded by a previous target stimulus, or by one or more warning signals or cues. Varieties of the RT task can be used to measure factors that influence general alertness and vigilant attention in animal models of cognitive performance in rats (Robbins, 2002), macaque monkeys (Baxter & Voytko, 1996), and also in humans ranging from healthy subjects to Alzheimer’s disease (AD) patients (McGuinness *et al.*, 2010). In these tasks RT – as a reliable measure of alertness – is known to be influenced by sleep deprivation (Dinges & Powell, 1985), pharmacological interventions (Unrug-Neervoort *et al.*, 1992; Killgore *et al.*, 2008) and age (Blatter *et al.*, 2006).

The effects of alertness on RT is particularly strong when the time interval between the pre-target warning cue and the target – the foreperiod – is relatively short and predictable, inducing rapid increases of phasic alertness in the expected moment of target appearance (Lawrence & Klein, 2013). However, if there is uncertainty in the timing of an expected event (e.g. random foreperiods from the 1-10 s duration range), the average RT is slower and attentional lapses occur more frequently for relatively short foreperiods (Tsunoda & Kakei, 2008; Basner & Dinges, 2011; Matthews *et al.*, 2017), whereas for long foreperiods, RT becomes faster (Klemmer, 1956; Niemi & Naatanen, 1981). Uncertainty in the timing of the imperative stimulus probably precludes the rapid burst of phasic alertness for early target arrivals (Hackley *et al.*, 2009; Weinbach & Henik, 2012). The way such uncertainty modulates RTs has been investigated using temporal attentional cueing experiments (Miniussi *et al.*, 1999; Coull *et al.*, 2000; Anderson & Sheinberg, 2008), where cues of varying validity predicted whether the target would arrive after a short or a long foreperiod. In those experiments, for short foreperiods, the valid cue resulted in shorter RT, and the invalid cue slowed down RT, while for long foreperiods, the validity of a cue had smaller or no effects on RT. These RT changes reflect the attentional costs and benefits of temporal orienting (Miniussi *et al.*, 1999; Coull *et al.*, 2000; Correa *et al.*, 2006; Lawrence & Klein, 2013). In the variable foreperiod scenario, the mere passing of time during the foreperiod induces continuously increasing subjective probability that the target would finally appear, aligning with the hazard rate, i.e. the objective conditional probability of appearance of the target given that it has not appeared yet. These probabilistic temporal expectations lead to changes in RT that are analogous to those observed in the case of explicit probabilistic temporal attentional cueing: the relative slowing of RTs after the shortest foreperiods – with low implicit, learned temporal expectations – corresponds to an attentional cost of the target appearing at an uncued time point – being unexpected based on explicit temporal expectations (Niemi & Naatanen, 1981).

Thus, besides alertness, temporal contingencies that are learned from the statistics of the environment induce temporal expectations, influencing RT in a way that is consistent with attentional effects measured using explicit cueing procedures (Nobre & van Ede, 2018). This in turn provides a simple and robust way to examine temporal expectation by measuring the foreperiod-dependence of simple RT (Milliken *et al.*, 2003; Weinbach & Henik, 2012). In human subjects, besides concomitant slowing of mean RT, foreperiod effects remain stable after repetitive testing (5-day practice, unpublished observations) or even after sleep deprivation (Tucker *et al.*, 2009) and are independent of the time of day, the prior duration of wakefulness and the relative fatigue occurring during task performance (time-on-task) (Matthews *et al.*, 2017), supporting the notion that foreperiod effects measure temporal expectation effects that can be dissociated from simple arousal or alertness-related factors.

In patients with cognitive decline, responses are generally slower and RT shows higher variability both within and between individuals compared to cognitively healthy elderly control subjects (Gorus *et al.*, 2008). Also, in contrast to young or healthy elderly subjects, AD patients show no RT improvement in the presence of alerting cues which predictably precede the target, indicating impairments in their ability to quickly and adaptively modulate phasic alertness (Tales *et al.*, 2002). Importantly, patients with AD express slower RT for early targets in a sequential auditory oddball paradigm compared to age-matched controls (Golob & Starr, 2000). Thus, besides the deficits of phasic alertness, deteriorated temporal expectation might also be a significant hallmark of pathological cognitive ageing, and assessing foreperiod-related effects in a simple RT task could prove to be a valuable method to dissect the impairments of temporal expectation within the framework of a highly translatable experimental paradigm.

Responding adaptively in the presence of temporal expectations is a complex process that involves basic sensorimotor neural circuits and higher-level networks of frontoparietal and striato-frontal loops and multiple neurotransmitter systems from the brainstem to the forebrain (Meck, 1996). For example, striatal dopaminergic lesion in rats abolished the foreperiod effect on the contralesional side by slowing down long-foreperiod responses (Brown & Robbins, 1991), while amphetamine improved short-foreperiod responding, also diminishing the overall foreperiod effects (Davis *et al.*, 2016) indicating the role of the dopaminergic system in time estimation and interval timing during temporal expectation. Complementary aspects of temporal expectation related to memory and attention have been frequently found to be linked to the cholinergic system (Meck, 2005). Especially, cholinergic neuromodulation has been found to be essential for behavioural adaptation to temporal contingencies (Parikh & Sarter, 2008; Howe *et al.*, 2010; Hasselmo & Sarter, 2011). In line with that, cholinergic deficits associated with pathological ageing are thought to be one of the most important mechanisms underlying impairments in cognitive function. Cholinergic modulations have been thoroughly investigated in reaction time tasks, establishing the role of the cholinergic system in alertness and attention in health and disease (Voytko *et al.*, 1994; Baxter & Voytko, 1996; Witte *et al.*, 1997; Howe *et al.*, 2010; Hasselmo & Sarter, 2011). However, studies of cholinergic effects on temporal expectation exploiting the variable foreperiod effect are surprisingly rare.

In the present study, therefore, to probe the influence of the cholinergic system in temporal expectation and alertness, we assessed foreperiod effects on RT and task performance in macaque monkeys. To fully exploit the information contained in RT data, we applied both parametric models and analyses that characterize the influence of treatments on the whole RT distribution (shift functions). Our first aim was to demonstrate the effects of foreperiod on RT and validate our paradigm for testing temporal expectation. Second, we investigated the effects of systemic muscarinic acetylcholine receptor suppression using scopolamine, a frequently used cholinolytic agent to induce transient amnesia in animal models of dementia (Ebert & Kirch, 1998). We also hypothesized that cholinergic inhibition, besides causing increases in RT and impairments in performance accuracy, would decrease the beneficial effect of foreperiod on RT, indicating a parallel impairment of temporal expectation. Finally, we looked at whether and to what extent would scopolamine-induced impairment of attentional functions be alleviated or reversed by the cholinesterase enzyme inhibitor donepezil, a cholinomimetic agent most commonly prescribed to alleviate symptoms in mild to moderate stages of AD.

## 2. Methods

### 2.1. Subjects

Five male 7 to 10 years old rhesus macaques (*Macaca mulatta*) were included in the study weighing 8 to 9 kg at the beginning of the experiments. One subject was excluded because of not displaying the foreperiod-dependent RTs we aimed to investigate (see details in the next section), therefore the final sample size was N=4. In order to ensure motivation during task performance, we applied a mild fluid restriction schedule on weekdays, where daily water intake, if not already consumed, was supplemented to 200 ml/day in the home cages. On weekends, water was available ad libitum. Animals were fed once per day, in the afternoons, following the daily testing session. Diet was standard nutritionally complete lab chow especially designed for non-human-primates (Altromin Spezialfutter GmbH, Lage, Germany) and was daily supplemented with fresh fruit and vegetable. In the home cage and testing rooms, temperature and humidity were maintained at 24 ± 1 °C and 55 ± 5 RH%, respectively.

All procedures were conducted in the Grastyán Translational Research Center of the University of Pécs. The study was approved by the Department of Animal Health and Food Control of the County Government Offices of the Ministry of Agriculture (BA02/2000-11/2012). Measures were taken to minimize pain and discomfort of the animals in accordance with the Directive 40/2013. (II.14.): ‘On animal experiments’ issued by the Government of Hungary, and the Directive 2010/63/EU ‘On the protection of animals used for scientific purposes’ issued by the European Parliament and the European Council.

### 2.2. Reaction time paradigm

Animals performed a simple RT task (Figure 1A) in one session per day. Each experimental session consisted of a total of 405 trials and lasted for approximately 60 min. During task performance, animals were seated in a primate chair in front of a computer screen (~50 cm distance). A response knob with an electric touch sensor was placed at a comfortable reaching distance (~20 cm) from the primate chair. At the beginning of each trial, a short tone was played to indicate that the subjects had to touch the response knob and prepare for the key release response. If the animal did not touch the knob within 2 s, the trial was not initiated and an intertrial interval of 2±0.5 s followed. If the sensor knob was touched, a warning stimulus (a black disc with a radius of 2.5 mm/ 0.57 diameter in degrees of visual angle in the centre of the screen) appeared within 0.3 s indicating the start of the foreperiod and remained displayed for the entire duration of the foreperiod. The duration of the foreperiod was set between 1.1 and 9.9 s (randomly drawn from 9 local probability bins: 1.1-1.9 s; 2.1-2.9 s; 3.1-3.9 s; 4.1-4.9 s; 5.1-5.9 s; 6.1-6.9 s; 7.1-7.9 s; 8.1-8.9 s; 9.1-9.9 s, incl. 45 trials per bin). When the foreperiod elapsed, the black disc turned white, which served as the target stimulus. The task required the animals to react as quickly as possible to the target stimulus by releasing the knob. In the absence of a response, the target stimulus disappeared after 1000 ms, the trial was considered unsuccessful and was terminated with no reward delivered. If the subject responded while the target was still displayed, the trial was completed correctly (completed trial). Upon a correct response the target stimulus disappeared, and subjects received a drop of liquid reward immediately after the response. If the trial was correct, then following a 2±0.05 s inter-trial interval the next trial started, if it was incorrect the inter-trial interval was 2±0.5 s. Stimulus presentation, response and reward delivery events were controlled by a script written by the authors in MATLAB programming environment (MathWorks, Natick, MA) using the Psychophysics Toolbox (Brainard, 1997; Pelli, 1997).

**Figure 1.**
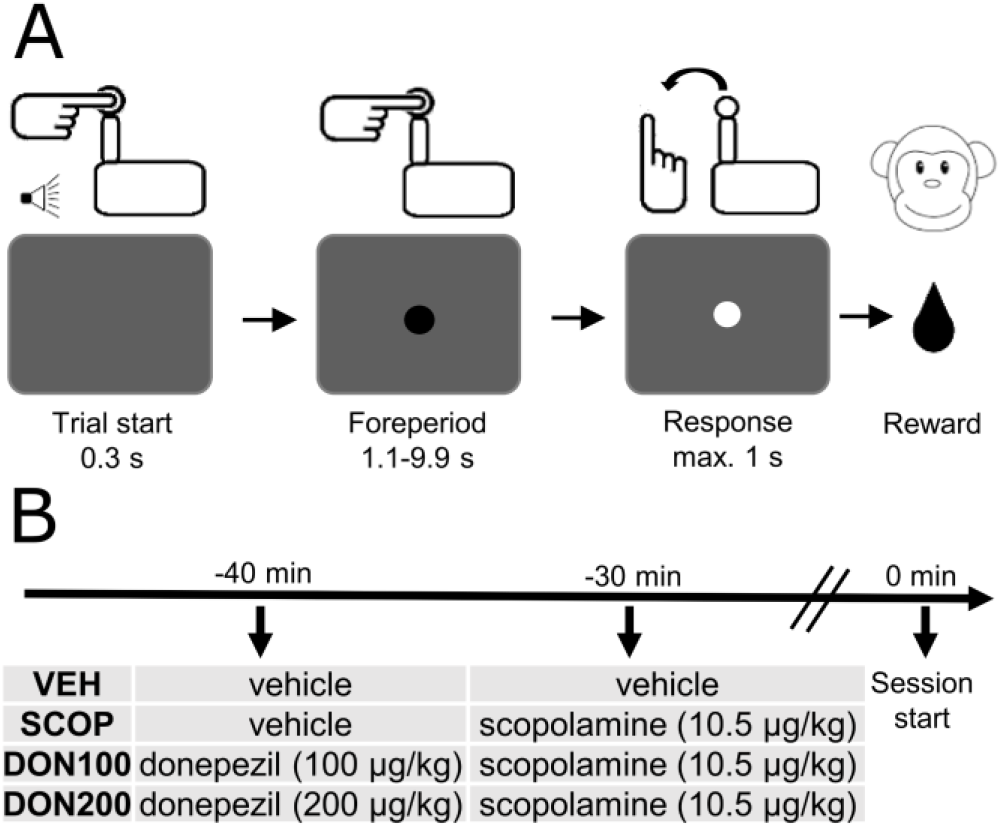
**(A) Schematic illustration of a single trial in the variable foreperiod simple RT paradigm.** On ‘Trial start’, animals had to touch a response knob after a short tone. Then, in the centre of the display, a warning stimulus (black disc) appeared and stayed on screen for a duration between 1.1 and 9.9 seconds, chosen randomly in each trial (Foreperiod). After the foreperiod the target stimulus (white disc) appeared. The task was to respond as quickly as possible to the target stimulus by releasing the knob (Response). If the subject responded in time the stimulus disappeared and the subject received the liquid reward. **(B) Schematic illustration of the timeline of treatments.** In each experimental session animals were administered two pharmacological agents (or vehicles). We administered the first injection at 40 and the second injection at 30 min before the task. We applied four types of treatment combinations, shown in the table under the timeline.

Animals performed the task at the same time of the day on every weekday. All animals had been previously trained to a criterion of at least 70% performance accuracy measured as performance rate (PR: number of correct (responded in time) trials divided by the number of initiated (touched the knob in time) trials and had been above criterion for at least five days consecutively prior to the start of the experiments.

In each subject, we modelled reaction time as a linear or quadratic function of the foreperiod, selecting the most appropriate model using the Akaike Information Criterion. In four out of five subjects, RT displayed a significant linear and/or quadratic dependence on foreperiod, indicating the presence of the temporal expectation that we aimed to investigate. One of the five subjects who were initially trained did not show temporal expectation in RT, therefore that subject was excluded from the study.

### 2.3. Procedures and drug administration

In the present placebo-controlled crossover and repeated measures experimental design, all subjects underwent 8 recording sessions with at least 72 hours of washout interval between the sessions. All experimental treatments were repeated twice. The first four sessions covered all the pharmacological treatment conditions, which were administered again for the next four treatment sessions; otherwise, treatment order was randomized and counterbalanced. To achieve stable and high plasma levels at the time of task performance, intramuscular injections of donepezil (Gedeon Richter Plc., Budapest, Hungary) were administered 40 min prior to behavioural testing, followed by scopolamine (Tocris Bioscience, Bristol, UK) or the corresponding vehicle treatment (saline) at 30 min before the behavioural testing session (Figure 1B). Donepezil and scopolamine were both dissolved in saline (saline, 0.9% NaCl). Saline was also used for vehicle (sham) treatments. Injection volume was set to 0.05 ml per kg bodyweight. There were four types of treatments (vehicle + vehicle (VEH); vehicle + 10.5 μg/kg dose of scopolamine (SCOP); 100 μg/kg dose of donepezil + 10.5 μg/kg scopolamine (DON00); 200 μg/kg dose of donepezil + 10.5 μg/kg dose of scopolamine (DON200)). Each solution was freshly prepared before each recording session and was stored for less than two hours.

Possible overt sedative side effects of scopolamine were subjectively observed by the experimenter. In initial scopolamine dose-finding pilot experiments, if the animal stopped responding during the task (which usually happened well before any overt sedative signs could be observed) we subsequently lowered the dose of scopolamine to preclude further overt sedative effects and to allow the desired continuous task performance. During the experiments, following a previously established rating scale (Arnsten *et al.*, 1988; Witte *et al.*, 1997) only normal or quiet arousal states were allowed (Scores 0 and 1) without drooping eyelids, slowed movements (Scores 2), intermittent sleeping (Scores 3), or complete sedation (Score 4).

### 2.4. Data analysis

In the analyses not involving foreperiod length as a factor or detailed analysis of RT distributions (see below), all sessions (8 per animal) of all animals (N=4) were analysed, leading to 32 sessions altogether. For the analyses of foreperiod dependence using a linear mixed model and RT distribution shift, the number of completed trials was insufficient in one scopolamine session of one animal, therefore that session was excluded from further RT analyses, leading to a dataset of 31 sessions from the 4 animals.

We analysed the ratio of completed trials (performance rate, PR), the number of omission and commission errors and RTs. Performance rate was determined by the number of correct (responded in time) trials divided by the number of initiated (touched the knob in time) trials. The number of trials with specific types of errors was also counted: A commission error occurred when the subject released the knob before the of appearance of the target stimulus, and an omission error occurred if an initiated trial ended without a response or with a late response (after the target disappeared). We also examined the effect of foreperiod on RT as the effect of temporal expectation. In the case of average PR, the effects of treatments and foreperiod were analysed with repeated measures of analysis of variance (ANOVA). To test sphericity we used Mauchly’s test and found that the sphericity assumption was not violated. In this analysis, Fisher’s Least Significant Difference (LSD) tests were used for post hoc comparisons. To test the number of commission and omission errors we used non-parametric related samples Friedman’s analysis of variance by ranks test. Here, Dunn’s test with Bonferroni correction was used for pairwise comparisons. These analyses were conducted using MATLAB (Mathworks, Natick, MA) and IBM SPSS (IBM Corp., Armonk, NY).

To test the interaction between the effect of treatment and foreperiod on RT, we used a linear mixed model on single-trial RT data. Foreperiod lengths were standardized for model fitting. Fixed effects were fitted for intercept, treatment, foreperiod and the treatment×foreperiod interaction, with treatment coded using Helmert contrasts (all non-VEH vs. VEH, DON100 and DON200 vs. SCOP, DON200 vs. DON100). Random intercepts for subjects were fitted, and sessions were nested within subjects with random terms for intercept and foreperiod. The model was fitted with the restricted maximum likelihood criterion using the lme4 (Bates *et al.*, 2015) package in R 3.6.0 (R Core Team, 2019) using the following model formula: `RT ~ 1 + treatment * foreperiod.z + (1 | subject.id) + (1 + foreperiod.z | subject.session.id)`, where `treatment` was a categorical factor with the 4 levels corresponding to pharmacological treatment conditions, `foreperiod.z` was the standardized foreperiod as a continuous variable, `subject.id` is the subject clustering variable for the 4 subjects and `subject.session.id` was the sessions-within-subjects clustering variable for the 31 sessions. P values were calculated using the Kenward-Roger approximation of degrees of freedom using the lmerTest package (Kuznetsova *et al.*, 2017).

Reaction time distributions are known to be non-normal and are theorized to contain more than one response component. To complement the parametric analysis and also capitalize on the rich information contained in the shape RT distributions, RT data was analysed by shift function analysis (Rousselet *et al.*, 2017) in MATLAB. The shift function describes the difference between the deciles of two distributions as a function of the deciles of one of the distributions. Deciles and medians are estimated using the Harrel-Davis quantile estimator in the shift function analyses and throughout the whole manuscript (including Figure 2). For statistical inference, confidence intervals of decile differences were estimated using the percentile bootstrap technique with n=10000 resamples (Wilcox *et al.*, 2014). First, we tested the differences of distribution of RT data between the SCOP and VEH, DON100 and SCOP, DON200 and SCOP, DON200 and DON100 sessions. Treatment sessions within the first four sessions and the four repetition sessions were compared separately; for example, data from the first SCOP session was compared to the first VEH session, and a separate shift function was used to compare the second SCOP session to the second VEH session. As an exception, for the animal with an omitted SCOP session, the remaining SCOP session was used in all relevant comparisons. Within each shift function analysis, bootstrap confidence intervals are adjusted for multiple testing across the 9 deciles. This method was used because a validated shift function estimation method encompassing the hierarchical structure of the present experimental design is currently not available – this aspect of inference is encompassed by the former linear mixed model analysis. Next, we analysed the difference between short (1.1-3.9 s) and long (7.1-9.9 s) foreperiod length categories within all the 31 treatment sessions included in the analysis (separately for both first and repeated sessions of VEH, SCOP, DON100, DON200).

**Figure 2.**
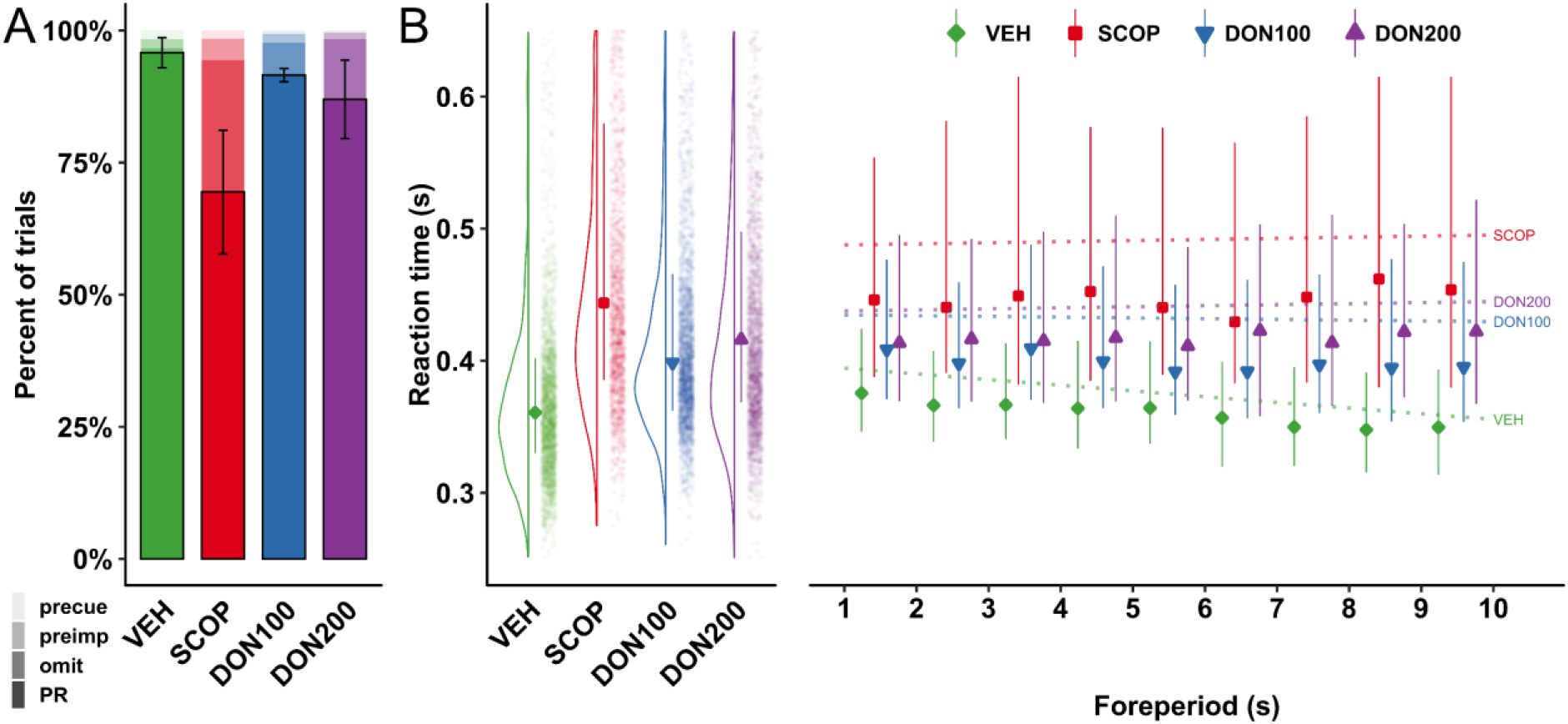
The effects of scopolamine and donepezil on mean performance rate (PR) and reaction time (RT). **(A)** Scopolamine (SCOP) significantly decreased PR compared to vehicle control (VEH), and this effect was partially reversed by donepezil (DON100, DON200). Bars with black contours filled with opaque colours show group mean PR, error bars here show s.e.m. Over the PR bar, areas shaded with decreasing saturation (legend in bottom right corner) show the proportion (among initiated trials) of omission errors (‘omit’), commission errors (‘preimp’) and responses preceding the warning stimulus (‘precue’). It is clearly seen here that the scopolamine-related impairment in PR is largely caused by omission errors. **(B)** The effect of scopolamine and donepezil on RTs and the foreperiod dependence of RTs for each of the four treatment combinations. On the left, RT distributions for each treatment are shown. The marker in the middle indicates the group mean of individual medians, the error bars cover the range from the 2^nd^ to the 9^th^ deciles (also group averages). The distributions of RTs pooled across subjects and sessions are also visualised with kernel smoothing (to the left) and as jittered dot plots (to the right, each dot is one trial). The right part of the plot shows RT for each 1-sec bin of foreperiod length, with markers and error bars using the conventions described above (group averaged 2^nd^, 5^th^ and 8^th^ deciles). Dotted lines correspond to the linear relationship between foreperiod as a continuous variable and RT as modelled by the linear mixed model in Section 3.2. Note that the distance between the marginal means of the linear model and the medians is commensurate with the skewness of the RT distribution. Reaction times decrease as foreperiod increases in the VEH treatment condition. Scopolamine treatment slowed down RTs and eliminated the effect of foreperiod on RT, and donepezil, while mitigating the overall RT increase, did not restore the foreperiod dependence in any of the doses applied.

## 3. Results

Animals showed reliable and high performance rates and short response latencies in the vehicle control sessions. Specifically, the average PR was 95.8 ± 2.8% (Mean ± s.e.m., see Figure 2A), the number of omission errors was 5 ± 1.2, and the number of commission (early response) errors was 13 ± 11.7. The group average of median RT was 361 ± 12 ms (see also Figure 2B). Reaction times were significantly shorter for longer foreperiods in vehicle sessions (linear mixed model, simple effect of foreperiod in VEH: t_34.7_=−3.31, p=0.0022; see Figure 2B; note that the presence of this pattern was an inclusion criterion for the study). However, foreperiod did not have a significant effect on PR (F_8,24_=1.59, p=0.18, η_p_^²^=0.35). Thus, we first analysed treatment effects PR without regard to foreperiod (Section 3.1), followed by analysis of RTs taking foreperiod-dependence into account as well (Section 3.2). Finally, using shift functions, we investigated how the effects of treatments (Section 3.3) and foreperiod (Section 3.4) varied across the whole RT distribution.

### 3.1. Analysis of task performance

The analysis of treatment effects showed a marginal main effect on PR (F_3,9_=3.28, p=0.073, η_p_^²^=0.52; see Figure 2A). In particular, scopolamine (SCOP) treatment significantly decreased PR compared to vehicle (SCOP vs. VEH post hoc p=0.017). Application of 100 μg/kg dose of donepezil (DON100) significantly reversed the scopolamine-induced impairments on PR (SCOP vs. DON100 post hoc p=0.037; see Figure 2A). 200 μg/kg dose of donepezil (DON200) had only a marginally significant effect on scopolamine-induced impairment on PR (SCOP vs. D200 post hoc p=0.085), however, there was no significant difference between the effects of the 100 and 200 μg/kg dose of donepezil (DON100 vs. DON200 post hoc p=0.623). These effects on PR were mainly driven by changes in the number of omission errors (failing to respond in time) rather than commission (false start) errors (Friedman’s ANOVA along the 4 treatments, omission: χ^2^(3)=10.8; p=0.013; commission: χ^2^(3)=3,9; p=0.272), with higher number of omissions under scopolamine treatment (157.3±71.8) compared to vehicle control (5±1.2; Dunn’s test for SCOP vs. VEH: p<0.01). The effect of donepezil on omissions relative to Scopolamine was not significant (DON100: 42.8 ± 9.5; DON200: 89.3 ± 61.2; Dunn’s test for DON100 vs. SCOP: p=0.6; Dunn’s test for DON200 vs. SCOP: p=0.6).

### 3.2. Reaction times and foreperiod effects

Linear mixed effects modelling confirmed a main effect of Treatment on RT (F_3,24.1_=15.2, p=10^−5^; see Figure 2A). Responses were significantly slowed down by scopolamine relative to control (contrast for all vs. VEH: t_24.0_=5.8, p=5×10^−6^), while donepezil partly reversed this impairment (contrast for DON100 and DON200 vs. SCOP: t_24.4_=−3.6, p=0.0015) in a dose-independent manner (contrast for DON200 vs. DON100: t_23.8_=0.55, p=0.59). Importantly, RT showed a clear and continuous dependence on foreperiod length in the VEH condition: RTs decreased as foreperiod length increased (simple effect of foreperiod in VEH: t_34.7_=−3.31, p=0.0022; see Figure 2B). Treatments clearly modulated this foreperiod dependence of RT (Treatment×Foreperiod: F_3,26.6_=3.29, p=0.036, Figure 2B): in contrast to control sessions, no such effect was observed in either of the treatment conditions (simple effects of foreperiod in all treatment conditions: |t|≤0.56, p≥0.58). This pattern is also reflected in an interaction contrast, wherein vehicle control sessions differed with respect to the foreperiod effect from all the treatment sessions (contrast for [all treatments vs. VEH]×Foreperiod: t_26.0_=3.06, p=0.0052). Treatment conditions did not differ from each other with regard to foreperiod effects (contrast for [DON100 and DON200 vs. SCOP] × Foreperiod: t_29.5_=−0.42, p=0.68, [DON200 vs. DON100] × Foreperiod: t_24.9_=0.74, p=0.47), and the main effect of delay was not significant (F_1,26.9_=1.47, p=0.24).

### 3.3. Effects of treatments on reaction time distributions

The effects of treatments on RT distributions are depicted in Figure 3A, where reaction times were pooled across sessions and animals, followed by kernel smoothing for visualisation purposes. To statistically characterize distribution effects, we analysed the treatment effects on RT in all deciles (1^st^-9^th^) using shift functions (Rousselet *et al.*, 2017) for pairs of sessions for each animal separately. Shift function analysis is a versatile tool for visualisation and inference, which entails estimating the deciles of the two distributions to compare and plotting their differences (see Methods and Figure 3B, C for details).

**Figure 3.**
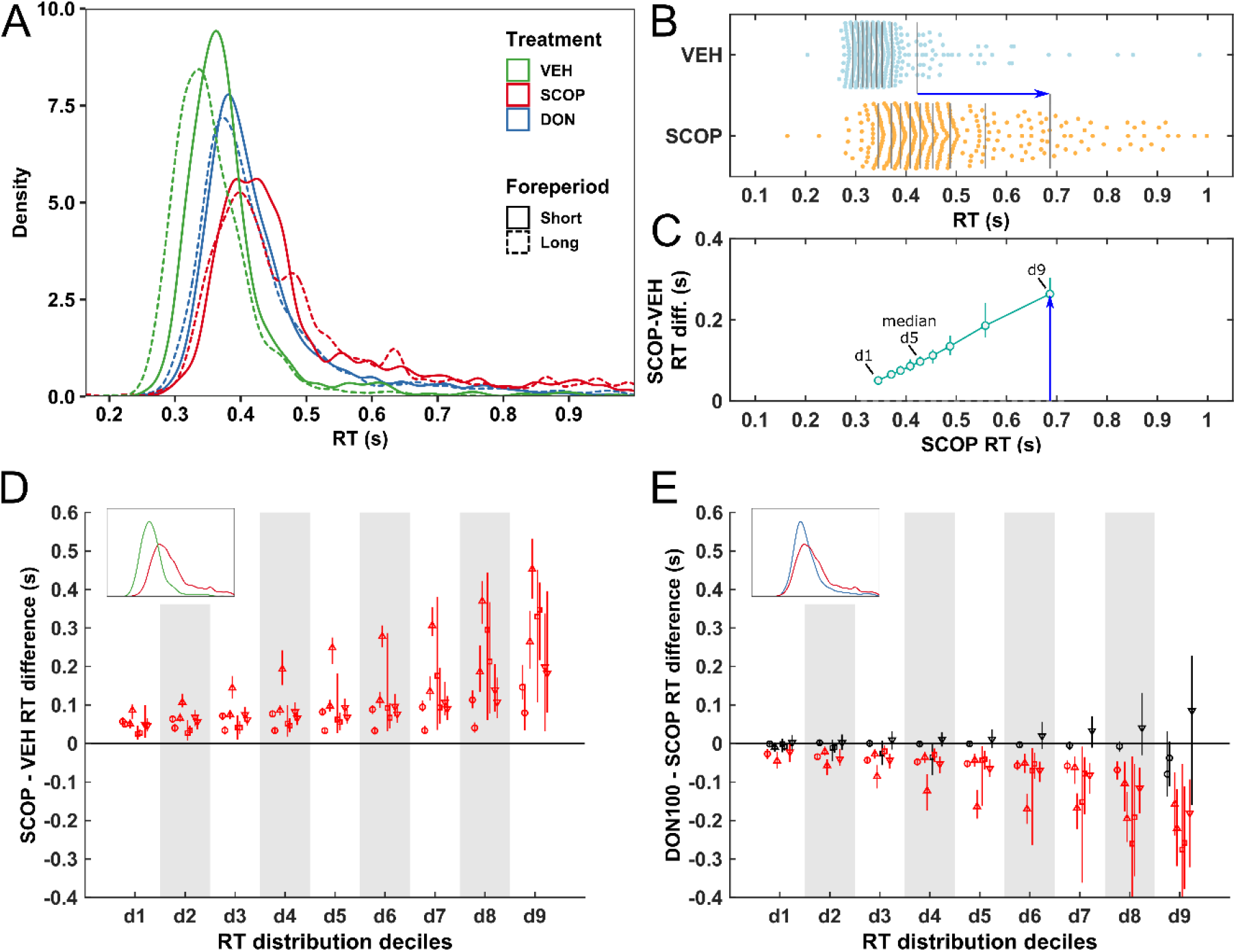
The effect of scopolamine and donepezil on RT distributions. **(A) Distribution of reaction times** in each treatment condition and foreperiod category (Short – continuous line: 1.1-3.9 s; Long – dashed line: 7.1-9.9 s), pooled across subjects and sessions. Data of the two donepezil doses was also pooled here as DON for visualisation to avoid clutter. Scopolamine treatment (SCOP, red curves) shifted the RT distribution rightwards (towards slower responses) compared to vehicle (VEH, green curves). This effect was partially compensated by donepezil treatment (DON, blue curves). Responses were slower for the short (continuous curves) compared the long foreperiods (dashed curves) in the vehicle control session (green curves), but this pattern is absent in all of the pharmacological treatment conditions. **(B) RT distributions from a VEH and a SCOP session** of one animal that illustrates the typical Scopolamine effect. Vertical grey lines indicate the deciles of the distributions. The blue arrow marks the shift of the 9^th^ decile in the SCOP session relative to its value in the VEH session. (**C) Shift function for the difference of RT distributions from the VEH and SCOP session shown above in***B*. In the shift function, the difference between SCOP and VEH at each decile is plotted against the decile values in the SCOP session. For example, the blue arrow denotes the same shift as in *B*, i.e. the large RT difference between SCOP and VEH in the 9^th^ decile. Vertical lines indicate the 95% simultaneous bootstrap confidence intervals. On the following panels, plots are composed of altogether 8 shift functions, 2 for each subject (see Methods). **(D) Shift functions depicting the effects of scopolamine treatment** relative to vehicle compared using shift functions. For each animal, the first SCOP session is compared with the first VEH session and the second SCOP session is compared with the second VEH session (see Methods). Marker shapes indicate individual animals. Here, instead of plotting against actual decile values as on *C*, deciles (labelled d1 to d9) are distributed evenly on the x-axes. The insets illustrate which distributions are compared using the colour key of Figure 3A. Red colour indicates significant differences. Scopolamine slowed down responses, especially at slower deciles. **(E) Shift functions comparing 100 μg/kg dose of donepezil**(DON100) with scopolamine treatment sessions, with the notation described above. Donepezil (DON100) decreases RTs relative to scopolamine monotreatment (SCOP), more effectively in slow deciles.

Scopolamine treatment increased RT relative to vehicle across the whole distribution (Figure 3A), but particularly in the case of slower RT, creating a strong tail of slow responses. This slow response component probably reflects attentional lapses. Shift analysis confirmed this, showing significant differences in all deciles between RTs in the vehicle and scopolamine treatment conditions for all sessions of all animals (Figure 3B, C, D). Application of donepezil significantly reversed the decreased RT (Figure 3A, blue curves, Figure 3E, Supplementary Figure 1A). In the case of the 100 μg/kg donepezil dose, the shift was significant across the whole distribution, except for one session of two animals (Figure 3E). Mirroring the effect of scopolamine, donepezil was more effective in the slower RT deciles, that is, it reduced the number of lapses. Reaction time distributions under donepezil treatment in the higher 200μg/kg dose were not significantly different from those of the lower dose, but visual inspection suggests that this dose might have been less effective (Supplementary Figure 1A, B).

### 3.4. The effects of foreperiod on reaction time distributions

To test the effect of foreperiod on RT distributions, we pooled foreperiods in two categories, short (1.1-3.9 s) and long (7.1-9.9 s), then compared them within each treatment using shift functions. In the VEH treatment longer foreperiods induced shorter RTs, especially in faster deciles (Figure 4A). Scopolamine treatment abolished the foreperiod dependence across the whole RT distribution (Figure 4B). Donepezil was not effective in any of the applied doses to alleviate the scopolamine-induced impairments on temporal expectation (Figure 4C, D).

**Figure 4.**
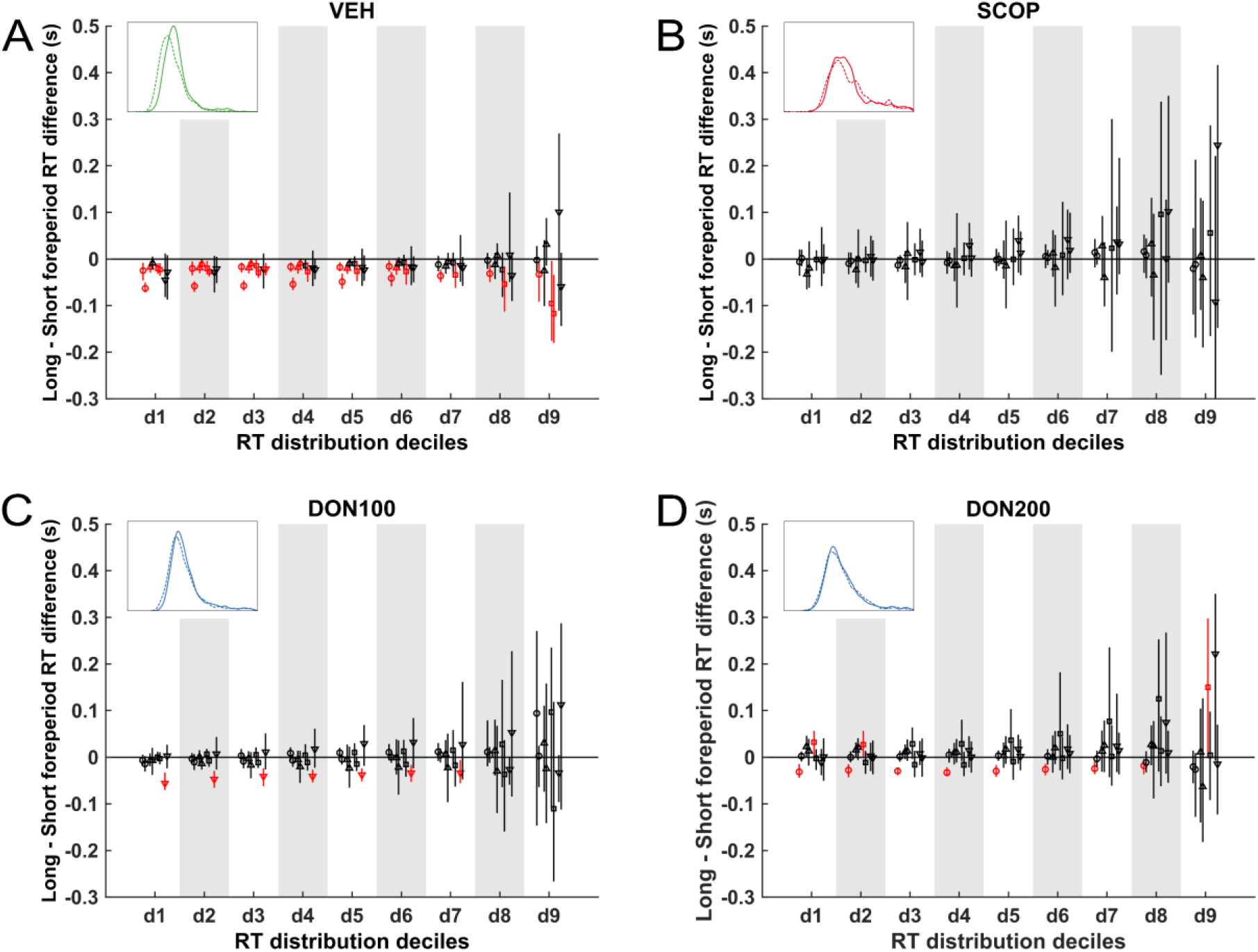
Shift functions showing the effect of foreperiod length (RTs from long minus short foreperiods) on reaction time distributions. Eight shift functions are plotted on each panel (except where noted), one for both sessions for each of the 4 animals. Insets illustrate which distributions are compared using the colour key of Figure 3A. Markers indicate individual animals and red colour indicates a significant shift. **(A)** For vehicle (VEH) treatment, longer foreperiods induced faster RTs, especially in the faster deciles. **(B)** Scopolamine (SCOP) treatment abolished the foreperiod effect. **(C, D)** During the donepezil treatments neither of the doses applied (DON100, DON200) were able to reverse the effects of scopolamine.

## 4. Discussion

In the present study, we applied a variable foreperiod simple RT paradigm to investigate the cholinergic modulation of temporal attention and alertness in macaques. The animals displayed a robust foreperiod effect in the vehicle control condition, i.e. responses were faster for longer foreperiods when temporal expectation (following the actual probability) of target appearance was strongest. The observed foreperiod effect primarily manifested in the faster deciles of the RT distribution, implying the presence of a fast response component (Noorani & Carpenter, 2016). Second, we went on to test how alertness and temporal expectation are modulated by the muscarinic acetylcholine receptor antagonist scopolamine as monotreatment or as co-administered with the cholinesterase inhibitor donepezil. Under scopolamine treatment, responses were generally slower, and also more lapses occurred, manifesting as a strong right tail component in the RT distribution (i.e., a shift and stretch of the RT distribution in the slow direction). In addition, the influence of foreperiod on RT was eliminated by the scopolamine treatment. Slower responses and lapses imply reduced alertness, whereas the lack of a foreperiod effect is a sign of impaired temporal attention. When donepezil was administered to alleviate scopolamine-induced impairments, task performance greatly improved and, though not reaching control levels, RTs became faster. Donepezil, mirroring the effects of scopolamine, was particularly effective at the slower deciles of the RT distribution, so it could largely alleviate the lapses introduced by scopolamine. In contrast to the alertness deficit, donepezil failed to mitigate impairments in temporal expectation, since the foreperiod effect was not reinstated under any of the donepezil treatment doses.

Although the general slowing of responses by scopolamine (Taffe *et al.*, 1999) and foreperiod effects have been demonstrated in a primate model (Witte *et al.*, 1996, 1997; Sharma *et al.*, 2015), the effects of cholinergic agents on temporal expectation in are much less investigated. Using a task that probes multiple attentional components in two macaque monkeys, Witte et al. (Witte *et al.*, 1997) showed reliable modulation of spatial, but not temporal attention by systemically administered nicotine. They also provided results on the effects of the muscarinic agonist atropine on temporal attention, but with rather ambiguous results (the effect increased with foreperiod in one animal and decreased in the other, see Table 2 in (Witte *et al.*, 1997)). Another previous study found that scopolamine caused a particularly strong response slowing and also strongly impaired spatial attentional orienting when locally infused into the intraparietal cortex, a key area of the attentional orienting network (Davidson & Marrocco, 2000). Although a foreperiod effect was found, pharmacological modulation of foreperiod effects was not tested, leaving open the question on the definitive role of intraparietal cholinergic activity in temporal attention. In our experiment, the foreperiod effect on RT was eliminated by the scopolamine treatment. This finding confirms that neuromodulation via muscarinic receptors plays an important role in temporal expectation.

We also investigated the effect of the acetylcholinesterase inhibitor donepezil on the scopolamine-induced deficit of temporal expectation and alertness. Donepezil is widely used for symptomatic treatment of AD (Martorana *et al.*, 2010), moderately improving the quality of life (Gauthier *et al.*, 2010) and several neuropsychological test results (Sugimoto *et al.*, 2002). Furthermore, cholinesterase inhibitors are known to be moderately effective in reversing scopolamine-induced cognitive and/or psychomotor impairments in humans (Wesnes *et al.*, 1991; Snyder *et al.*, 2005) and macaques (Buccafusco *et al.*, 2008). Here we confirmed that donepezil partially reversed scopolamine-induced slowing in a simple reaction time task; however, it failed to attenuate the scopolamine-induced impairment on temporal expectation, since the foreperiod effect was not restored. We put forward that this is because donepezil may improve general alertness but not temporal expectation, which only together may serve as the functional basis of optimized task performance.

Multiple lines of evidence support that temporal expectation can facilitate neural processes from several stages of sensory processing (Ghose & Maunsell, 2002; Fischer *et al.*, 2013; Wiegand *et al.*, 2017) through higher-level attentional processes (Seibold & Rolke, 2014) up to response selection and execution in motor and premotor areas (Doherty *et al.*, 2005; Cui *et al.*, 2009; Pomper *et al.*, 2015). From the range of candidate brain areas and mechanisms that could underlie the cholinergic modulations observed in our results, the prefrontal cortex (PFC) merits further discussion, as in this area the link between cholinergic activation and temporal attention is the best described. In particular, Parikh et al (2007) established the role of prefrontal cholinergic transients in the processing of a warning stimulus in a rodent model (see also Parikh & Sarter, 2008; Hasselmo & Sarter, 2011). They have shown that the time course of the cholinergic transients evoked by warning stimuli depends on the length of the cue-to-reward interval (foreperiod), peaking right before the expected time of reward delivery. Also, before successfully detected warning stimuli, cholinergic activity in the medial PFC was found to show a negative trend, conversely, missed trials were preceded by gradually increasing cholinergic activity. Thus, in general, the temporal dynamics of cholinergic activity appears to follow the expected time course of events with relevance to perception (warning stimulus) or action (reward or imperative stimulus), playing a crucial role in the expectation of behaviourally relevant stimuli, and also during stimulus-driven shifts of attention. Given that the role of prefrontal areas in temporal expectation is also well established in humans (Vallesi *et al.*, 2007), suppressing cholinergic activity in the PFC is a promising candidate mechanism for scopolamine-induced impairments of temporal attention observed in our study. Also, considering the strong RT modulation effect of scopolamine in the intraparietal area together with the notion that the activity of neurons in the intraparietal area are known to display robust signals of temporal expectation (Janssen & Shadlen, 2005), it is reasonable to suggest that the intraparietal area is also a probable site of action where scopolamine can modulate temporal expectation.

A possible neuropharmacological mechanism that could underlie the present results is that muscarinic receptors might play a role in either the production of cholinergic transients or the readout (Howe *et al.*, 2017) of their temporal dynamics, and these mechanisms could be required for adaptive behavioural responses to temporal contingencies in the environment (Parikh *et al.*, 2007; Parikh & Sarter, 2008; Hasselmo & Sarter, 2011). Further, it can be conceived that the increased availability of ACh under donepezil treatment can partly restore the cholinergic tone and thus tonic behavioural alertness, but at the same time, acetylcholinesterase activity would be required for the temporal specificity of phasic prefrontal cholinergic activity, for example by facilitating the decay phase of cholinergic transients. Under high tonic ACh levels resulting from inhibited acetylcholinesterase activity, cholinergic transients might be smeared, leading to lower signal-to-noise ratio of the temporal attentional processes they normally support. In line with this, research on rats suggests that the temporal specificity of cholinergic transients during sustained attention, and concomitant better task performance can be achieved by targeting specific receptor subtypes (Howe *et al.*, 2010) – pharmacological modulation of tonic ACh levels probably cannot achieve this (Trocme *et al.*, 2010). This raises the possibility that pro-cholinergic drugs that increase the sensitivity of particular subtypes of ACh receptors, i.e. agonists, and especially positive allosteric modulators, might be potentially more successful in supporting higher level cognitive functions that require the adaptive dynamics of phasic cholinergic neuromodulation.

Interpretation of the results should also take into account the complex relationship between noradrenergic and cholinergic systems. Indeed, the locus coeruleus (LC) noradrenergic plays a role in selective attention functions. Low performance due to insufficient attentional resource allocation has been associated with low LC activity (Aston-Jones *et al.*, 1994). In addition, noradrenergic afferents derived from the LC have been shown to control working memory and attentional functions located in the dorsolateral PFC (dlPFC) (Ramos & Arnsten, 2007; Arnsten, 2011), where phasic LC activity correlated with focused or selective attention, and tonic LC activity correlated with flexibility and scanning during task performance (Aston-Jones *et al.*, 1999). Moreover, noradrenaline levels in the dlPFC were reportedly elevated in the case of unexpected uncertainty (e.g., attention set shifting) while acetylcholine levels were elevated in the case of expected uncertainty (e.g., during foreperiods preceding rule-based responses) (cf. review article by Yu & Dayan, 2005). Together, the two transmitter systems may have a synergistic and also permissive role in shaping the appropriate behavioural response through acting in the dlPFC and elsewhere to fine tune higher order cognitive control during task performance. As it is known that scopolamine influences the activity of intrinsic LC neurons (Egan & North, 1985; Adams & Foote, 1988) by antagonizing muscarinic acetylcholine receptors, it is reasonable to suppose that attentional resource allocation in the midbrain may, at least partly, also be jeopardized when scopolamine is used (mainly as an amnestic agent).

Reaction time distributions are known to be non-normal and composed of multiple response components, therefore conventional parametric models do not fully capture potential treatment effects. To this end, we analysed treatment effects across the whole RT distribution using a nonparametric quantile-based method. The foreperiod effect was more expressed in the fast RT deciles, and this effect was eliminated by scopolamine. This suggests that temporal expectation is chiefly manifested in a fast response component that might require attentional and motor preparatory processes involving cholinergic mechanisms on muscarinic receptors. Fast response components are also associated with ‘urgency’, elevated readiness to respond and lower criterion values in decision theoretic models (Reddi & Carpenter, 2000; Noorani & Carpenter, 2016). This is adaptive in a simple RT task like ours, but it is also known that responses speeded up by the foreperiod effect are less accurate in tasks involving more complex cognitive control mechanisms (Correa *et al.*, 2010; Weinbach & Henik, 2013; Korolczuk *et al.*, 2018).

In sharp contrast to foreperiod effects, the effect of both scopolamine and donepezil was stronger on the slower tail of the RT distribution. While donepezil mitigated the response slowing caused by scopolamine to some degree, the fast response component for long foreperiods did not reappear under any of the donepezil treatment doses. Considering the slow right tail component thickened by scopolamine and partly mitigated by donepezil, in the framework of psychomotor vigilance tasks these responses are usually considered lapses of attention, i.e. temporary disengagement of sustained attention from task performance (Basner & Dinges, 2011). We think that this interpretation holds best in the case of our study; the interesting alternative explanation that the faster responses may be impulsive and inaccurate (Correa *et al.*, 2010; Korolczuk *et al.*, 2018) and, conversely, the slow responses may be more accurate, cannot be proved or disproved in the framework of the currently used simple RT paradigm. Conjointly, the dissociation of foreperiod and pharmacological effects across the two tails of the RT distribution also supports our proposition that scopolamine caused impairments in temporal attention that donepezil failed to restore, along with effects on alertness that could partly be mitigated.

To characterize the effect of the muscarinic antagonist scopolamine and cholinesterase inhibitor donepezil on temporal expectation and alertness, here we measured the foreperiod effect in a variable foreperiod simple RT task in six macaques. Scopolamine decreased response speed, which was interpreted as a reduction in alertness that could partly be recovered by donepezil. In addition, scopolamine was shown to abolish temporal expectation (as indexed by the foreperiod effect), supposedly through impairing cholinergic activity evoked by warning stimuli that is known to optimize attentional performance. We also showed that pharmacologically increasing the availability of ACh by donepezil is, on its own, not sufficient to reverse the observed deficits in temporal expectation. Analyses of RT distributions provide further support for our conclusions by showing that temporal expectation is manifested in fast responses, while cholinergic pharmacological effects were more prominent on slow responses most probably caused by attentional lapses due to reduced alertness. The present results also help delineate the efficacy and scope of donepezil and other cholinomimetic agents as cognitive enhancers in present and future clinical practice.

## Acknowledgements

The authors would like to thank Judit Inkeller for valuable technical contribution. The authors also thank Anna Káldi, Zoltán Gödri and Bence Petrovai for their assistance in animal care. This work was supported by Gedeon Richter Plc. and the Hungarian National Brain Research Program of the National Research, Development and Innovation Office of the Hungarian Government (grant number ‘2017-1.2.1-NKP-2017-00002’) and the Hungarian Higher Education Programme for Excellence (Felsőoktatási Intézményi Kiválósági Program, FIKP) [20765-3/2018/FEKUTSTRAT]). The funding bodies did not influence the study design, the collection, analysis and interpretation of data and the decision to submit the article for publication.

## Author Contributions

V.O., A.T., B.L. and I.H. designed the research. V.O. conducted the experiments, V.O., A.T. and B.K. performed data analysis. V.O., B.K., B.L. and I.H. wrote and reviewed the manuscript.

## Data Accessibility Statement

The datasets generated during and/or analysed during the current study are available from the corresponding author on reasonable request.

## Abbreviations

ACh: Acetylcholine
AD: Alzheimer’s disease
LC: Locus coeruleus
PR: performance rate
RT: Reaction time
NHP: Non-human primate
SCOP: scopolamine
DON: donepezil

