## Supplementary Material for "Dissociating cholinergic influence on alertness and temporal attention in primates in a simple reaction time paradigm"

***Corresponding Author:**

Grastyán Translational Research Center, University of Pécs

Mailing address: 6 Ifjúság út, H-7624, Pécs, Hungary

**Supplementary Material**

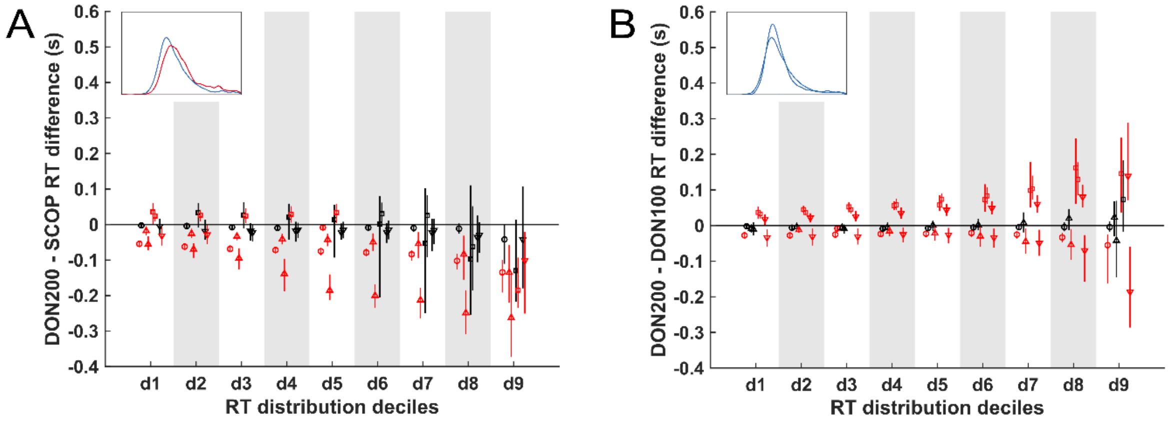


**Figure S1 (A) Effects of 200 µg/kg dose of donepezil (DON200)** compared with scopolamine (SCOP) in shift function for each session of each animal separately. The insets illustrate which distributions are compared using the colour key of Figure 3A. Markers indicate individual animals. Red colour indicates significant differences. The scopolamine-induced impairment of reaction time (RT) was partially reversed by donepezil. **(B) Shift functions comparing 100 µg/kg dose of donepezil (DON100) with 200 µg/kg dose of donepezil (DON200).** In half of the experimental sessions DON200 was more effective than DON100 and in the other half of the experimental sessions DON100 was more effective. Because of this, RT distributions under DON100 were on average not different from DON200.
